# From Past to Future: The Impact of Climate Change on a Mediterranean Lizard

**DOI:** 10.64898/2026.03.26.714419

**Authors:** Arda Cem Kuyucu

## Abstract

Mediterranean Basin, one of the most important hot spots for reptiles, is also expected to experience significant impacts with climate change, posing a severe risk for the herpetofauna of the region. This study uses the snake-eyed lizard *Ophisops elegans* as a model organism to investigate the potential impacts of past and future climate change on reptile distributions in the region. An ecological niche model (ENM) was developed with the Maxent algorithm, with location points from GBIF and bioclimatic variables from the CHELSA dataset, then projected onto past LGM (∼21 kya) and future (2071-2100 SSP3-7.0 and SSP5-8.5) scenarios. Results show that the present-day distribution of *O. elegans* is primarily driven by temperature seasonality and precipitation, indicating a preference for coastal Mediterranean climates with dry summers. The LGM projection suggests a fragmented and contracted range, confined to coastal refugia around the Mediterranean and Caspian Seas. Future projections for 2071-2100 show consistent and alarming contraction of suitable habitats under both SSP scenarios. In conclusion these findings indicate that *O. elegans* is vulnerable to significant habitat loss under projected climate change. This severe impact on a wide-spread species implies that the herpetofauna of the Mediterranean Basin may face a significant threat in future.

## Introduction

Ecological niche models (ENMs) are powerful tools that quantify species-environment relationships to map potential habitat suitability. By projecting these models onto past (e.g., Last Glacial Maximum, LGM) and future climates, ENMs provide vital insights for conservation, as historical responses to climate shifts can inform predictions of future changes (Whittaker et al. 2005; Yousefi et al. 2015; Maguire et al. 2015; Rej and Joyner 2018; Walkup et al. 2022; Feng et al. 2024). Understanding the past responses is crucial, as species that experienced range contractions during previous climatic periods may exhibit similar shifts in response to future changes (Araújo et al. 2008; Sillero and Carretero 2013). Furthermore, developments in ENMs have advanced our understanding of how species might respond to future climate change (Maguire et al. 2015).

As ectotherms, reptiles are highly dependent on external temperature and precipitation, making them susceptible to climate change-induced distributional shifts and extinctions, particularly given their limited dispersal ability (Sinervo et al. 2010; Bonino et al. 2015; Sanchooli 2017; Nowakowski et al. 2018; Tan et al. 2023). The Mediterranean basin, a hotspot for both reptile diversity and climate change, is experiencing significant warming, decreased precipitation, and increased weather extremes that are already causing biodiversity declines. (Hoerling et al. 2012; Sternberg et al. 2015; Grillakis 2019; Albano et al. 2021; Aurelle et al. 2022; Zittis et al. 2022). ENMs can aid in identifying risks to species, determining key conservation areas, and helping to build viable conservation strategies for the distinct herpetofauna of this region (Barbosa et al. 2012; Srinivasulu et al. 2021; Biancolini et al. 2025).

The genus *Ophisops* is a very wide spread genus that is distributed in a large area that span North Africa, Middle East and South Asia (Agarwal and Ramakrishnan 2017; Montgelard et al. 2020). Among these, the snake-eyed lizard, *Ophisops elegans*, is a common lacertid distributed across the Eastern Mediterranean and Iran (Kyriazi et al. 2008; Oraie et al. 2014; Montgelard et al. 2020). The ancestral lineage of *O. elegans* originated in Southwest Asia, and its expansion into the Middle East and Eastern Mediterranean is assumed to be connected to successive climatic shifts toward cooling and aridification, which promoted open habitat expansion during the Late Miocene and contributed to the formation of the Sahara Desert (Zhang et al. 2014; Montgelard et al. 2020). Suggesting that the distribution of *O.elegans* is determined by its sensitivity to humidity and the preference for open habitats which are connected with precipitation levels. Additionally lizard physiology is directly connected with ambient temperature and their performance is very sensitive to extreme temperatures (Clusella-Trullas et al. 2011; Morley et al. 2019), therefore changes in minimum and maximum temperatures are expected to be crucial determinants of the distribution of *O. elegans*. Despite the fact that the formation of all clades occurred long before LGM, continuing patterns of shifts to cool and arid conditions linked to the expansion of high-latitude ice sheets might have caused vicariance events reinforcing the allopatric distribution of clades. Despite the severity of LGM conditions certain areas might also have been refuge areas by maintaining suitable microclimate conditions for lizards. The global sea level drop during the LGM might have resulted with an expansion of coastal lowlands in certain regions sutiable for temperate species (Marske et al. 2009), it is shown that these expanded coastal areas became refuges for other Mediterranean lizards *Podarcis wagleriana* and *Podarcis siculus* (Senczuk et al. 2017; 2019). Ecological niche modelling approaches can be instrumental in determining hot spot regions for any potential intra-specific units, considering that Southern Anatolia, Levant and Iranian plateau compose an important key region of reptile biodiversity according to the reptile biodiversity maps prepared by Roll et al. (2017). Choosing a widespread species like *O. elegans* as a model organism has some key advantages; firstly ENMs built with a larger number of locations across varied environments tend to have lower sampling bias and attain higher accuracies (Qiao et al. 2015, 2017). Second, widespread species’ higher distribution and adaptation to diverse environments increases the degree of transferability to past and future projections as models built on rare species are more prone to overfitting (Breiner et al. 2015; Della Rocca et al. 2019).

This study aims to 1) identify the bioclimatic drivers of the current distribution of *O. elegans*, hypothesizing that minimum temperature and precipitation are critical limiting factors; 2) reconstruct its LGM distribution, expecting a more reduced and fragmented range; and 3) project its future (2071-2100) distribution, anticipating habitat contraction due to increased warming and aridity. To answer these questions, bioclimatic variables from the CHELSA database and *O. elegans* occurrence records from the literature were used to build ecological niche models with Maxent, projecting climatically suitable areas for past, present, and future scenarios.

## Material and Methods

### Study Area and Occurrence Records

The main distribution area of *Ophisops elegans* is the coast of Mediterranean from Western Anatolia to the coast of Syria and Lebanon with pockets of distribution from inner Anatolia to Iranian Plateau, in the East Mediterranean and Middle East (EMME) region. The region exhibits strong climatic trends: with North EMME being mostly temperate with dry and warm to hot summers and relatively mild to very cold temperatures (especially in mountainous regions) while the south is marked by large, arid and hot regions with sparse vegetation (Lelieveld et al. 2012; Zittis et al. 2022). Average minimum temperatures range from -18 □ in mountainous areas to 5 □ in coastal regions, while average maximum temperatures can reach 40-50 □ in desert areas (Kostopoulou et al. 2014). Precipitation also exhibits very high variation, while some areas such Taurus and Caucasus mountains have up to 2000 mm/year precipitation, in the south extremely arid areas have near zero precipitation (Francis et al., 2021). This vegetation covers relatively diverse types of habitats from Mediterranean type coniferous forests and shrublands to steppe vegetation of Anatolia and Iran and deciduous forests in the northern regions near Balkans, Black Sea and Caucasus (Tüfekcioğlu and Tavşanoğlu 2022; Ugarković et al. 2022).

Occurrence records for *O. elegans* were gathered from the Global Biodiversity Information Facility (GBIF). Records were filtered to remove points with location uncertainty greater than 1 km and those outside the study rasters. To reduce sampling bias, the points were spatially thinned to 5 km using the spThin R package (Aiello□Lammens et al. 2015), resulting in a final dataset of 433 occurrences (Fig. 1). These filtered occurrences are available in supplementary information.

**Fig. 1.**
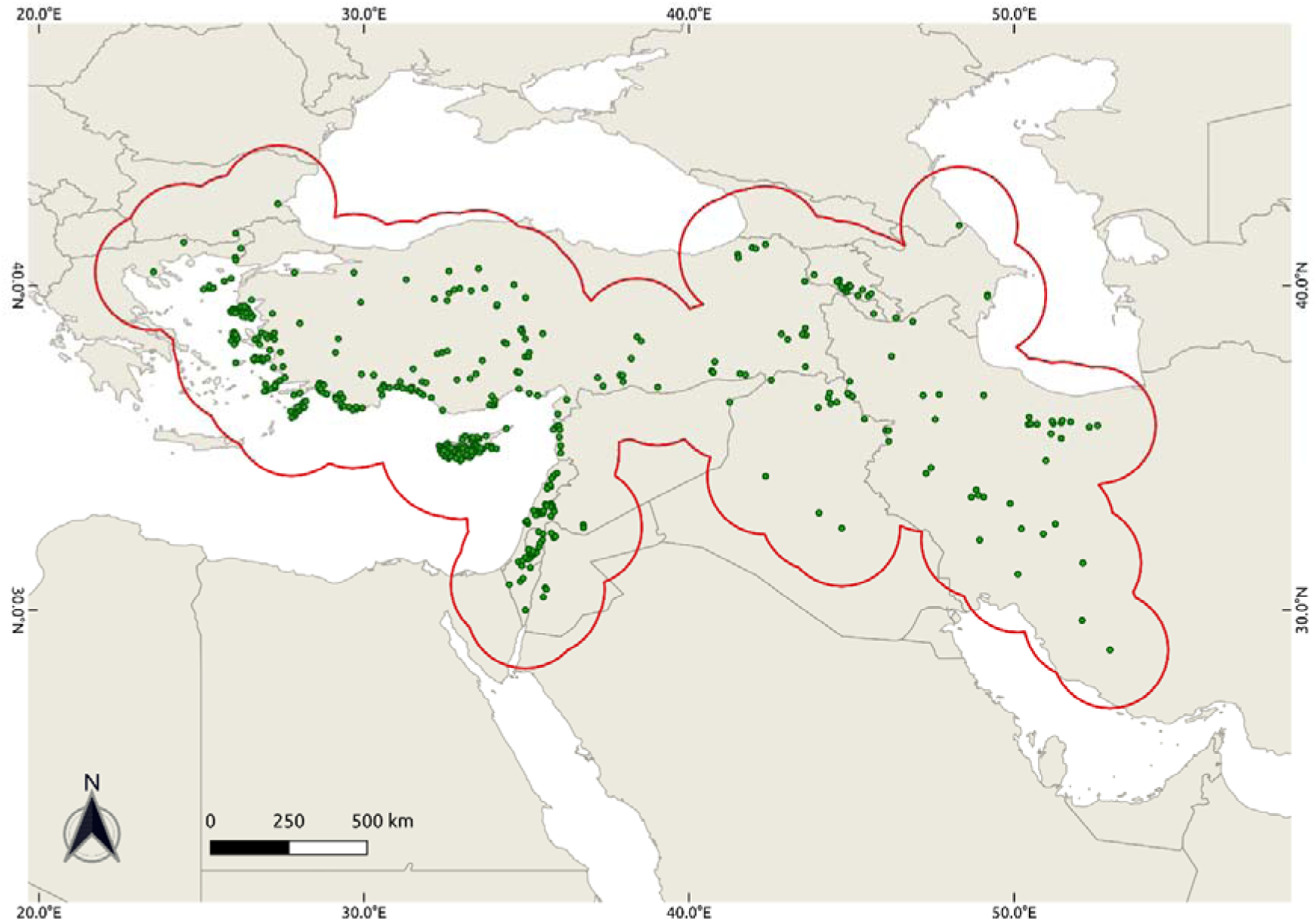
*Ophisops elegans* locations used in building the ecological niche models after 5 kilometer thinning and filtering. Red lines show the circular point buffer area used to build the models.

### Environmental Variables

Variables for the ecological niche model and projections were gathered from the CHELSA dataset available at https://chelsa-climate.org/exchelsa-extended-bioclim (Karger et al. 2023) with a 30 arcsec, (∼1 km) resolution. To build the distribution model, climatic variables for the 1981–2010 period were used. Bioclimatic factors 8, 9, 18 and 19 were not used as there are inconsistencies related to spatial artefacts which are first observed on the Worldclim dataset but these are also present in the CHELSA data (Escobar et al. 2014; Aguilar-Domínguez et al. 2021; 2022; Escobar 2020). Environmental rasters were clipped to the area of interest between latitudes 19° and 49° and longitudes 15° and 57° WGS84. All geographic computations were performed with QGIS (QGIS Geographic Information System 2022), the GDAL library in Python (Open Source Geospatial Foundation 2022), and R version 4.2.2 (R Core Team 2022).

### Future and Past Projections

All GCM scenarios used in this study are shown in Table 1 with explanations. For the future and past projections, data that are based on the implementation of the CHELSA algorithm on “Coupled Model Intercomparison Project” CMIP6 (Future) and “Paleoclimate Modelling Intercomparison Project Phase 3” PMIP3 (Past LGM) data (Karger et al. 2017) were used. For the Last Glacial Maximum (∼21 kya), we used CHELSA LGM data based on the implementation of the (PMIP3), which is a specialized sub-project that serves as the paleoclimate component of CMIP5 (Braconnot et al. 2012) (available at https://chelsa-climate.org/paleo-climate/). Bioclimatic parameters were downloaded from CHELSA dataset for 5 selected PMIP3 scenarios which include MRI-CGCM3 (Yukimoto et al. 2012), MPI-ESM-P (Jungclaus et al. 2011), MIROC-ESM (Sueyoshi et al. 2013), IPSL-CM5A-LR (Sepulchre et al. 2019) and NCAR-CCSM4 (Gent et al. 2011). For the future projections, GCM’s available for CMIP6 2071-2100 of SSP3-7.0, a midway scenario and SSP5-8.5, the worst-case scenario, were used for future projections (available at https://chelsa-climate.org/cmip6/). These include UKESM1-0-LL (Sellar et al., 2019), MRI-ESM2-0 (Oshima et al. 2020), MPI-ESM1-2 (Mauritsen et al. 2019), IPSL-CM6-LR (Boucher et al. 2020) and GFDL-ESM4 (Dunne et al. 2020).

**Table 1.**
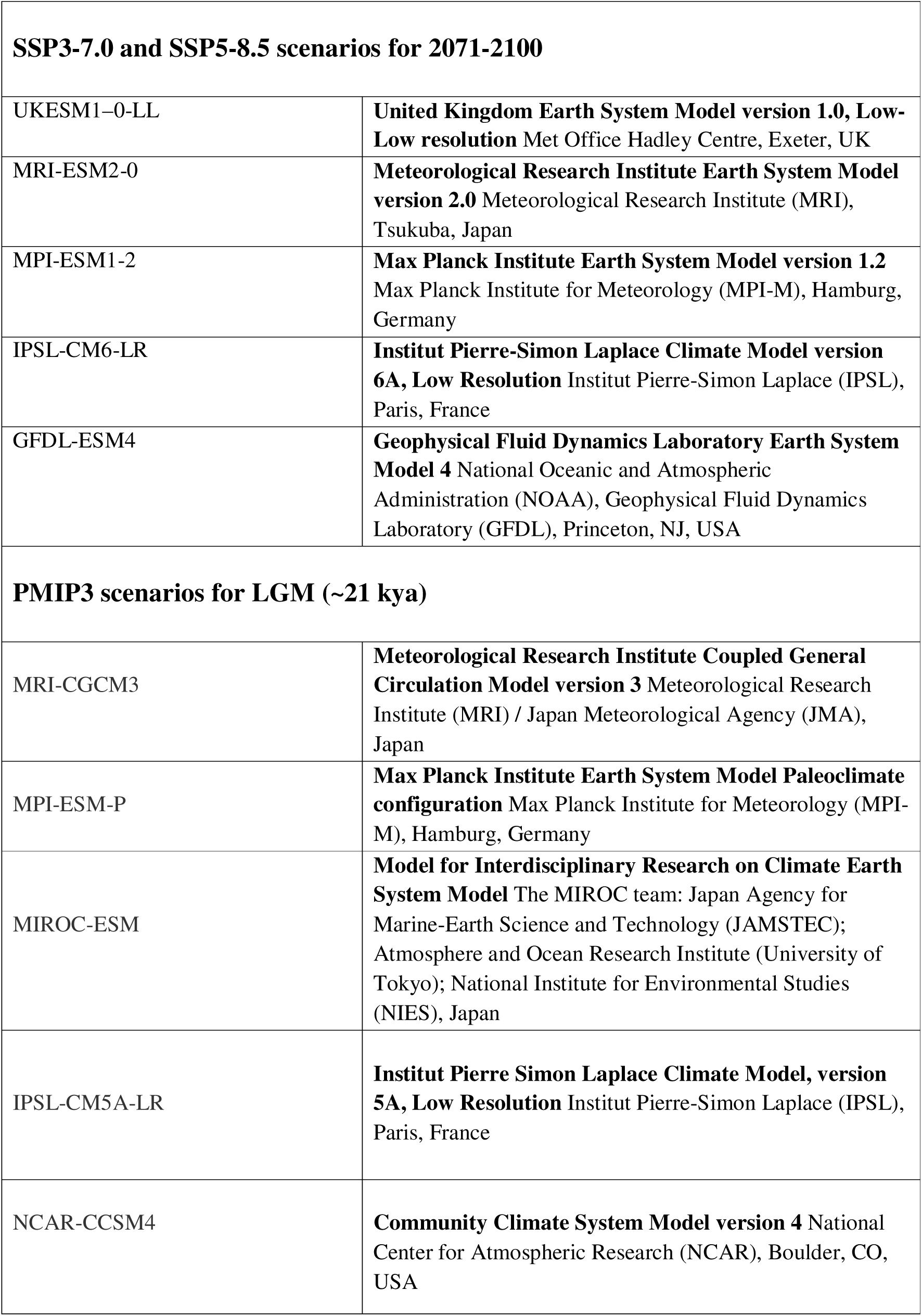
All GCM scenarios used in model projections.

### Ecological Niche Models

Locations were split into a training set (n = 273, ∼60%), a cross-validation test set for preliminary evaluations (n = 118, ∼30%) and an independent secondary test set for final evaluation (n = 45, ∼10%). Depending on the Grinnelian niche concept, ENM’s require an M space that depicts the accessible area for species (Soberón 2007; Barve et al. 2011; Soberón and Peterson 2020). To define this M space, we created a 200 km buffer around sampling points using the “ellipsenm” package for R, available at https://github.com/marlonecobos/ellipsenm (Cobos et al. 2020). 200 km was chosen as a buffer since it was the minimum buffer radius that provided a connected M space (Fig. 1). Variable selection was done by using the vifcor function in the usdm package for R (Naimi 2017), choosing variables from correlated pairs (0.8 threshold) depending on the variance inflation factor (VIF), and also by consulting the previous studies on lizards, special focus was given to parameters related with seasonality, maximum and minimum temperatures and precipitation (Oraie et al. 2014; Giacometti et al. 2024). The set of bioclimatic parameters used to build the models are shown in Table 2. Distribution models were created with the maximum entropy algorithm (Phillips et al. 2006) using Maxent version 3.4.4 (Steven et al. 2021) embedded in the Kuenm package for R (Cobos et al. 2019). 250 candidate models were created with 5 combinations of linear, quadratic, product and hinge features (L, LQ, H, LQH, LQPH) and 50 regularization parameters between 0.1 and 5 with a step of 0.1 and 10,000 random background points were used. Maximum thresholds was set as 0.05 for the omission rate, while a minimal threshold of 0.7 was chosen for area under curve (AUC) and a pROC (partial receiver operating characteristic) test was used according to Peterson and Soberón (2012) and Aguilar-Domínguez et al. (2021). The final model was selected from among the candidates conforming to the above criteria using the Akaike Information Criterion corrected for small sample sizes (AICc), with a maximal ΔAICc criteria of 2 (Nuñez-Penichet et al. 2021; Warren and Seifert 2011). The final model was tested with the independent test set that was set aside before model calibrations, and model projections were built with 10 bootstrap replicates. To account for the effects of model projections under novel future conditions, models were transferred with allowed projections in novel climate conditions (extrapolation with clamping) (Cobos et al. 2019).

**Table 2.** All bioclimatic variables used in the model with percent contributions.

| Variable | Percent Contribution |
| --- | --- |
| Bio4 (temperature seasonality) | 52.4 |
| Bio14 (precipitation amount of the driest month) | 20.1 |
| Bio13 (precipitation amount of the wettest month) | 12.6 |
| Bio15 (precipitation seasonality) | 7.2 |
| Bio6 (min. air temp. of the coldest month) | 3.8 |
| Bio5 (max. temperature of the warmest month ) | 2.1 |
| Bio3 (isothermality) | 1.8 |

Suitability maps were created by dividing the logistic suitability maps into five categories. 10^th^ percentile training presence logistic threshold was set as the first threshold then followed by category bins of 25%, 50% and 75% suitability classes. For future projections, a consensus map was calculated by taking the median of the 5 GCM scenarios for SSP3-7.0 2071-2100 and 5 GCM scenarios for SSP5-8.5 2071-2100 and another consensus map for past LGM was created by taking the median of all 5 LGM PMIP3 scenarios. The uncertainty of the suitability model for the current conditions was mapped using the standard deviation of the 10 replications of the model, while for the GCM projections uncertainties were calculated by taking the standard deviation of 5 different scenarios separately for PMIP LGM and SSP3-7.0 2071-2100 and SSP5-8.5 2071-2100 models. To determine the degree of extremity of novel past and future conditions a Mobility Oriented Pair (MOP) analysis was carried out and extrapolation risk of transfer regions was calculated with nearest 10% reference for each GCM projection (Owens et al. 2013; Flores-López et al. 2022).

## Results

### Model Parameters

A total of 250 candidate models were created for evaluation. All these candidate models were statistically significant, 119 met the 0.05 omission rate criteria and only one met the ΔAICc criteria of ≤ 2. This final model is an LQPH model with a regularization parameter of 2.3 (mean AUC ratio = 1.59, omission rate = 0.035, ΔAICc = 0, pROC ≤ 0.001). The final evaluations done on the independent test set also met the aimed criteria with an omission rate of 0.048. The percentage contribution of all 7 predictor variables can be seen in Table 2. The most important environmental variable was Bio4, temperature seasonality with a contribution of 52.4%, followed by Bio14, precipitation amount of the driest month with 20.1% contribution, these are followed by Bio13 (12.6%) and Bio15 (7.2%), unexpectedly minimum temperature was behind these with only 3.8% contribution.

### Past, Current and Future Predictions

All suitability predictions are shown in Fig. 2. 10^th^ percentile training presence logistic threshold was 0.13. The climatic suitability belt stretches from Aegean islands and Coast to Levant and North Africa and to Iran with the highest suitability concentrated on the Mediterranean Coast. The LGM distribution shows a markedly reduced suitable area, concentrated around the Mediterranean and Caspian coasts, with a prominent gap between these two regions. Additionally, the overall degree of suitability is lower, with almost no area showing suitability higher than 0.75 under LGM conditions. SSP3-7.0 2071-2100 projections show a significant reduction especially in inner regions including Central Anatolia and Iran, additionally in the coastal areas highly suitable regions are almost non-existent except very small patches in Southwest Anatolia and Levant. Interestingly, SSP5-8.5 is very similar to SSP3-7.0, only with very small additional reductions.

**Fig. 2.**
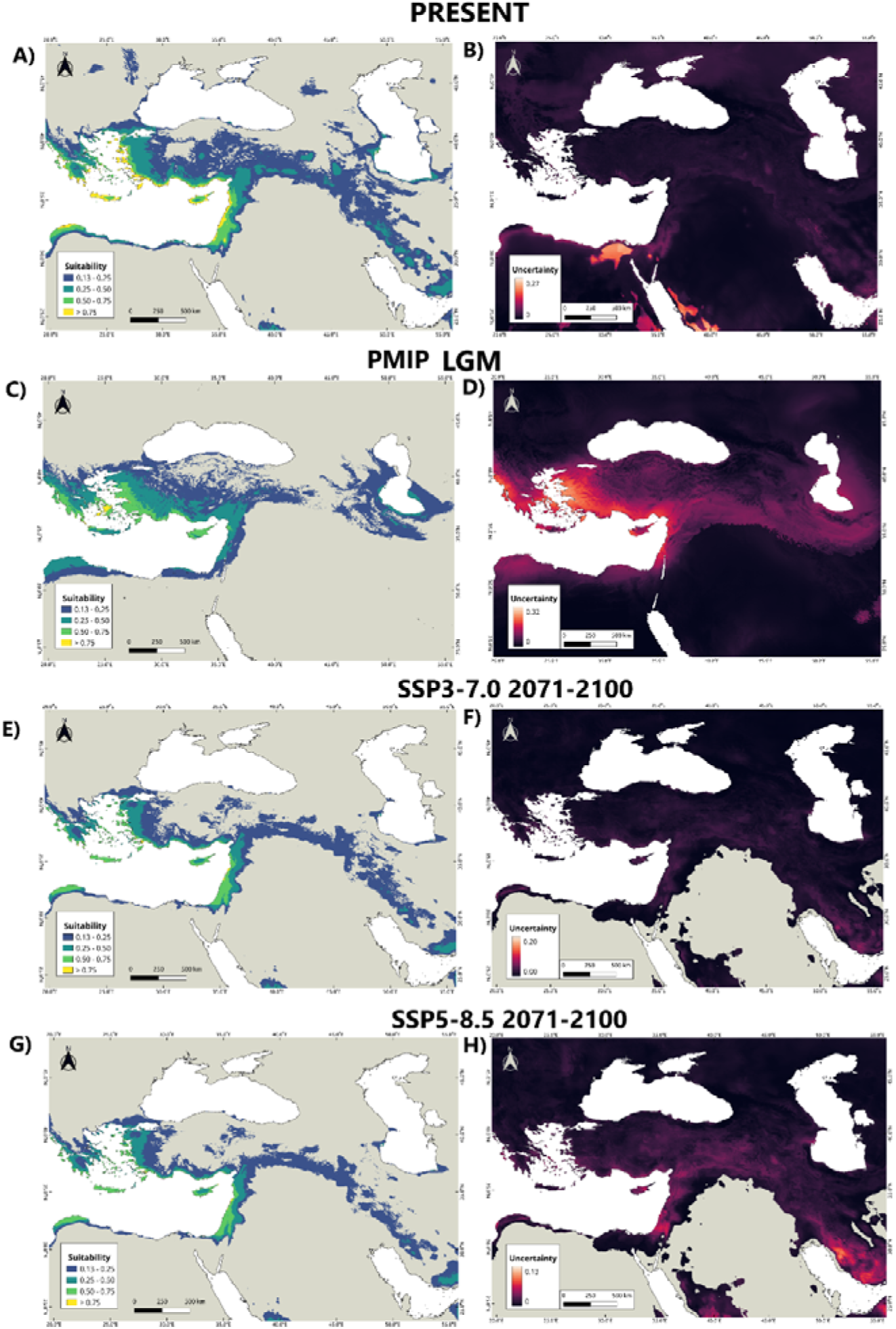
Side by side suitability and uncertainty values for all projections, 0.13 is the 10^th^ percentile training presence logistic threshold. **A)** Suitability for the present conditions **B)** Uncertainty for the present conditions. **C)** Suitability for the PMIP LGM conditions calculated as the average of 5 GCM scenarios. **D)** Uncertainty for the PMIP LGM conditions. **E)** Suitability for the SSP3-7.0 2071-2100 conditions. **F)** Uncertainty for the SSP3-7.0 2071-2100 conditions. **G)** Suitability for the SSP5-8.5 2071-2100 conditions. **H)** Uncertainty for the SSP5-8.5 2071-2100 conditions.

### 3.3 Uncertainty and MOP Results

Uncertainty results for the current conditions indicate that the degree of uncertainty is negligible in the main distribution areas. LGM results on the other hand show that the level of uncertainty is very high in the main distribution areas according to the LGM projections, showing that PMIP3 LGM scenarios differ greatly from each other. MOP results also show that due to the much more extreme conditions of LGM all areas show some degree of extrapolation, with the areas of high extrapolation being in the North and South regions while middle regions including Anatolia and Caspian Sea show the lowest degree of extrapolation. The future SSP3-7.0 and SSP5-8.5 scenarios did not show any significant extrapolation risk within the species’ projected distribution, as almost all areas of high extrapolation risk were located outside of the suitable regions. Uncertainties among different GCM scenarios are also very low for both SSP3-7.0 and SSP5-8.5. All MOP results for individual scenarios are included in the supplementary information.

## Discussion

### Current Distribution and Environmental Parameters

The predicted present-day suitability aligns well with the known distribution of *O. elegans*, concentrating on coastal Mediterranean habitats characterized by hot, dry summers and wetter non-activity periods. Precipitation during non-activity periods likely boosts prey availability in the subsequent dry season (Dendi et al. 2019; He et al. 2025). The results also showed that, contrary to our expectation, minimum temperature was not as important compared to precipitation and seasonality with only a 3.8% contribution. This may be because diurnal lizards can use thermally buffered refuges to avoid extreme cold (Grenot et al. 2000; Wilson and Cooke 2004; Berman et al. 2016), or because its effect is masked by the stronger influence of seasonality and precipitation variables in the model.

### Past and Future Projections

As expected, LGM projections showed a much more restricted distribution, with suitable habitats confined to coastal refugia around the Mediterranean and Caspian seas. This reflects the extremely dry and cold steppe conditions that dominated the region during that period (Carolin et al. 2019; Davis et al. 2024). Additionally, in the LGM projections the suitable areas are clustered around the Mediterranean and Caspian sea where the coastal areas are larger due to the reduction of sea levels, previously studies on the wall lizards in Italy and fungus beetles in New Zealand projected wider distributions in the emerged coastal lowlands in the LGM (Marske et al. 2009; Senczuk et al. 2017; 2019). The South Caspian coast, in particular, is known to have been an important faunal refuge (Parvizi et al. 2018; SaberiLPirooz et al. 2021; Stadelmaier et al. 2025). However, these interpretations must be treated with caution, as our MOP analysis confirmed high extrapolation risk and high uncertainty among GCMs, highlighting that model choice significantly impacts LGM predictions

As can be seen from the map the suitable areas during LGM are concentrated around three regions: the East Mediterranean coast from Greece, West and Inner West Anatolia to the South to Levant, the coast of the Caspian Sea and North Africa. Studies showed that the past climatic events during Miocene climatic fluctuations and Plio-Pleistocene glacial cycles was one of main causes of formation of numerous reptile clades (Paulo et al. 2008). These regions show significant gaps between each other during the LGM and although clades have formed much before this period it is highly probable that earlier similar events during Miocene and Pleistocene were instrumental in the formation of these clades (Kyriazi et al. 2008; Montgelard et al. 2020). Additionally, although the clade formation occurred earlier, fragmentations during the LGM might have contributed to the reinforcement of the already formed clades. All in all, we should also be careful before interpreting these projection results, because LGM conditions generally represent very extreme conditions compared to the current, pointing to high degrees of extrapolation and uncertainty. In the present study for the LGM all the areas had some degree of extrapolation with the main distribution areas for *O. elegans* having lower degrees compared to the more extreme parts in the North and South. This is inevitable as there is not yet any way to eliminate extrapolation in the model transfers, especially in projections to the much more distant past (Varela et al. 2015; Guevara et al. 2019). Additionally, different GCM scenarios show significant differences between each other which can also be seen in uncertainty maps. Thus our results also reinforce the notion and results of the previous models that the type of GCM chosen has a significant impact on the distribution predictions in LGM (Collevatti et al. 2013; Varela et al. 2015;Chefaoui and Serrão 2017; Guevara et al. 2019).

Both future GCM projections SSP5-8.5 and SSP3-7.0 for 2071-2100 show a reduced area of climatic suitability. Most of the decline seems to occur in inland habitats but there is a decrease in the degree of suitability in all areas including coastal regions. In regard to uncertainty, future projections are much more reliable compared to LGM projections, this is no wonder that they project a much closer time, 45-75 years in future compared to the 20,000 years past. On the other hand, the reduction in climatic suitability is calamitous compared to this relatively short time change. These dramatic changes and shifts in distribution are already predicted by many models and studies, in fact we are already experiencing these changes in abundance and distribution of reptiles (Bowler et al. 2017; Lenoir et al. 2020; Biber et al. 2023). As ectotherms reptiles face greater risk since they are much more sensitive to environmental changes and additionally many species are distributed in regions where habitat temperatures are close to their thermal tolerance limits (Sinervo et al. 2010; Piantoni et al. 2016; Diele-Viegas and Rocha 2018; Morley et al. 2019).

We must also take into consideration the uncertainty as one of the main challenges in this regard is the disagreement between the different GCM’s on the state of climate in future (Wright et al. 2015) on the other hand uncertainty and extrapolation parameters are much more lower for future projections compared to the LGM. Although the climatic projections are relatively more reliable there are other complicated factors. For example, projections from correlative ecological niche models might not capture other potential risks such as the effect of extreme weather events as bioclimatic variables generally include parameters that are summed over months or seasons like means of temperature and precipitation or seasonality. The magnitude, duration and frequency of extreme events are expected to increase with climate change and in fact even now we are experiencing a dramatic increase in these events, challenging the adaptation ability of many reptiles species (Kyselý et al. 2012; Liz et al. 2019; Morley et al. 2019; Sinervo et al. 2024). One of these risks concerns the potential climate traps during late winter and early spring such as early emergence which increases the risk of exposure to sudden freezing temperatures following the warm spells (Turner and Maclean, 2022). In this regard in addition to the correlative niche models such as employed in this study, the integration of mechanistic models that use functional physiological traits that depend on microclimatic parameters holds importance (Amano 2012; Kearney et al. 2010, 2018; Martins et al. 2025). In conclusion, although *O.elegans* does not seem to be at risk of extinction in the future, the projected contraction in suitability is alarming for a widely adapted species and would indicate much more endangering conditions for species with more restricted distribution such as endemic species or species that are already under risk of extinction.

## Acknowledgments

The numerical calculations reported in this paper were performed with TUBITAK ULAKBIM, High Performance and Grid Computing Center (TRUBA resources).

## Funding

This research did not receive any funding

## Data Availability

Location points used in this study are available in the supplementary information at https://data.mendeley.com/z99nmfrzhx.

## Conflicting interests

The author declares no conflicting interests.

## Notes

https://data.mendeley.com/z99nmfrzhx.

### Competing Interest Statement

The authors have declared no competing interest.

