## supplementary information for "From Past to Future: The Impact of Climate Change on a Mediterranean Lizard"

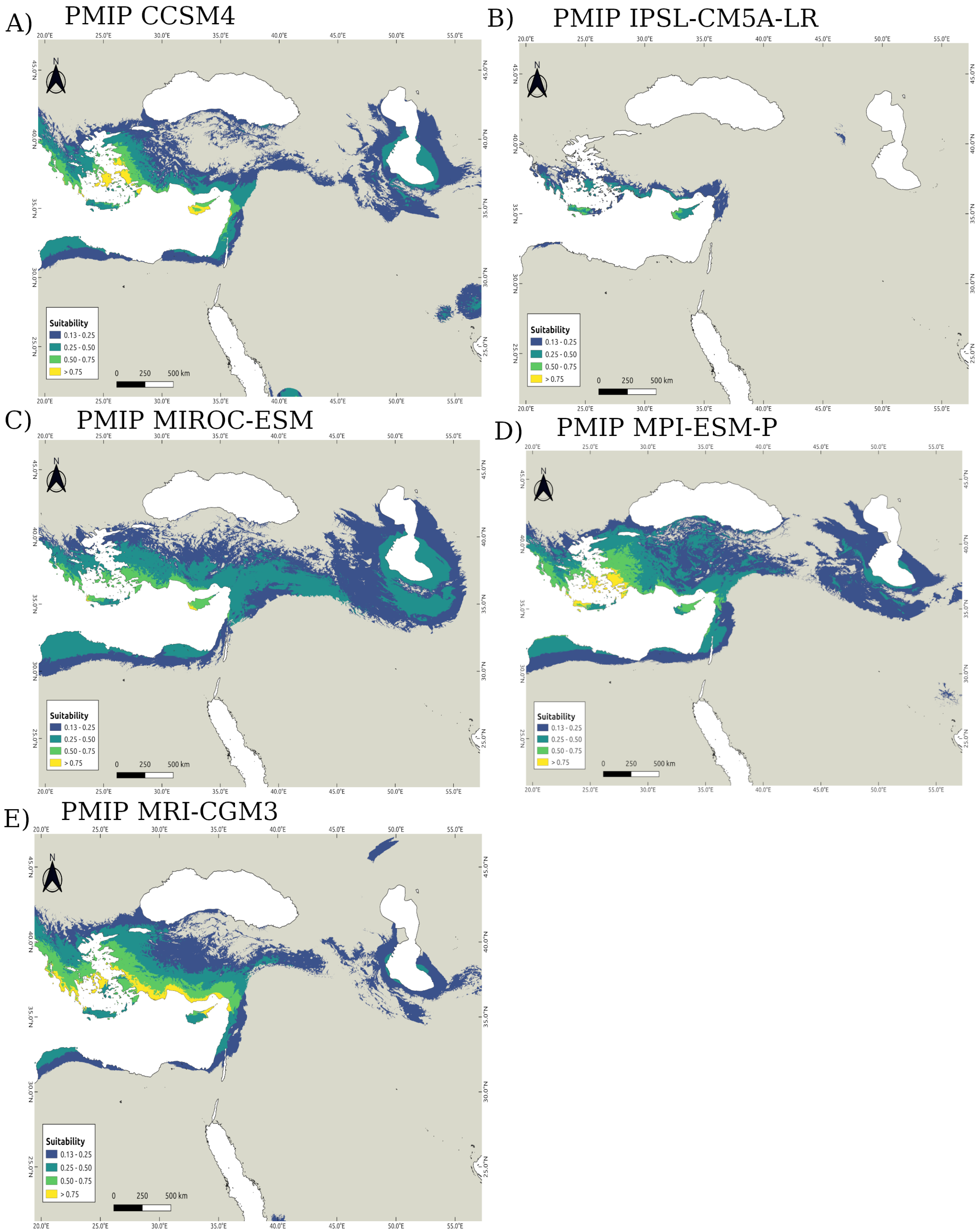


Suitability maps for all LGM PMIP GCM scenarios

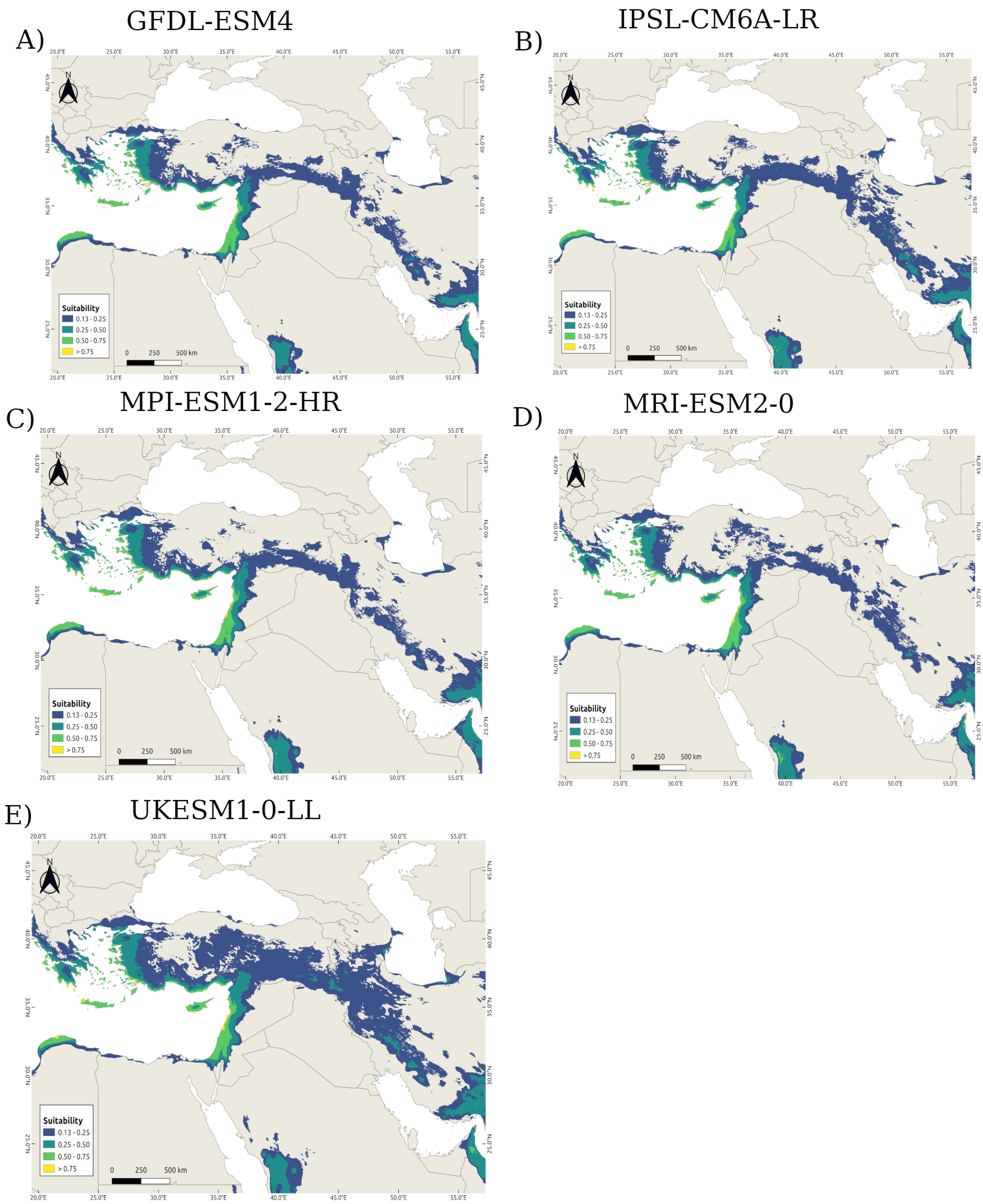


Suitability maps for all SSP3-7.0 2071-2100 Scenarios

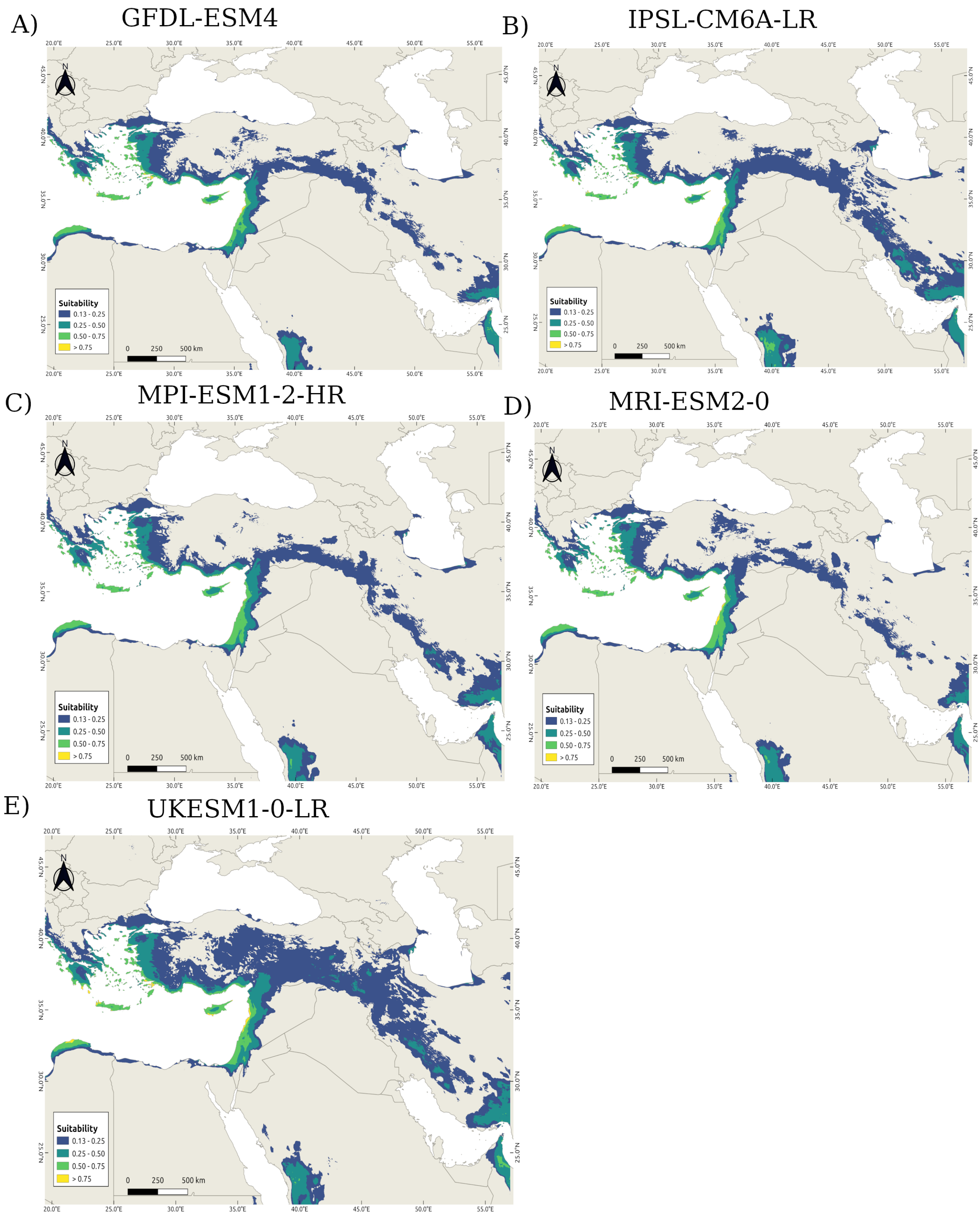


Suitability maps for all SSP5-8.5 2071-2100 scenarios

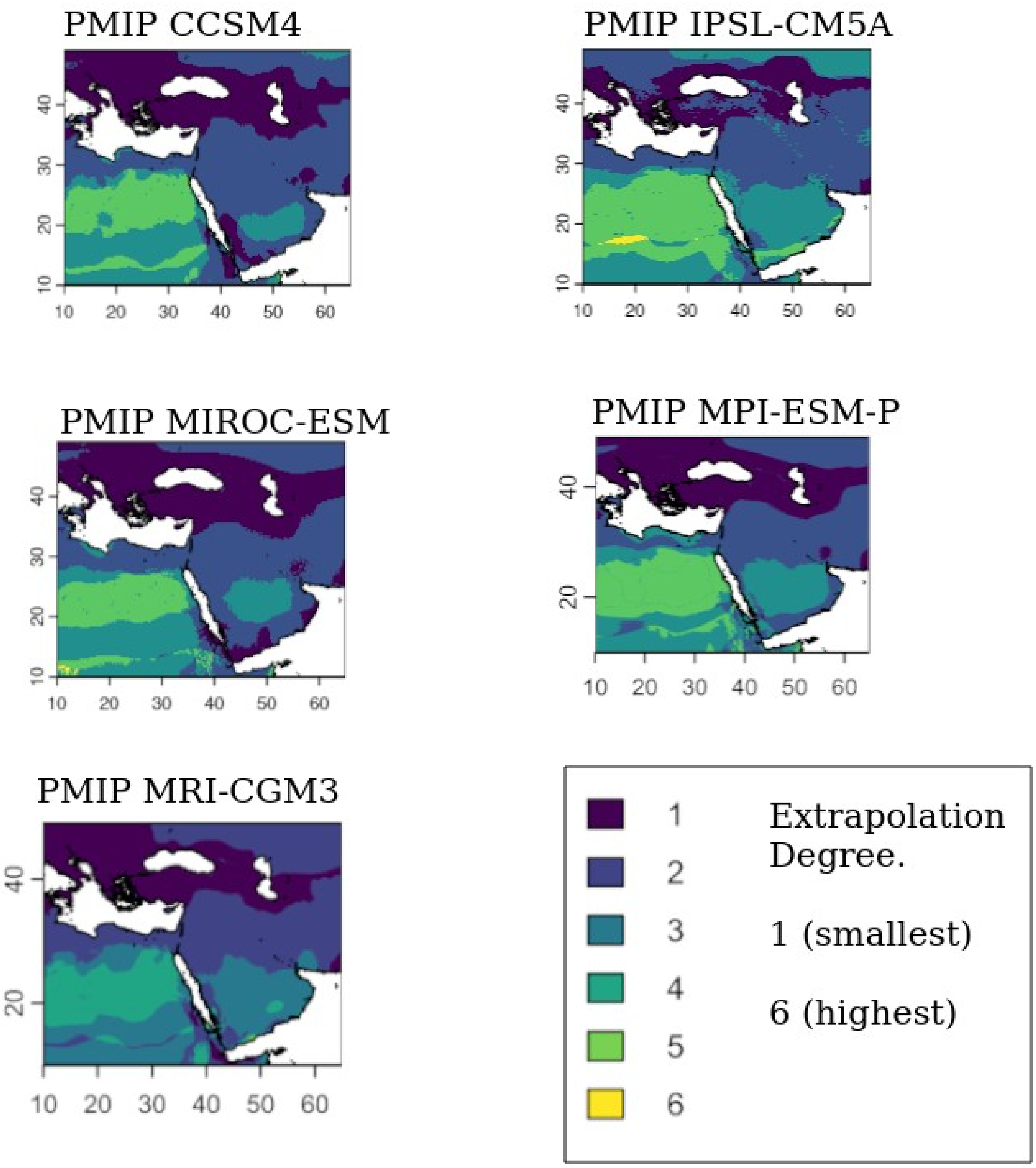


MOP (Mobility Oriented Pair Analysis) extrapolation maps for all PMIP LGM scenarios.

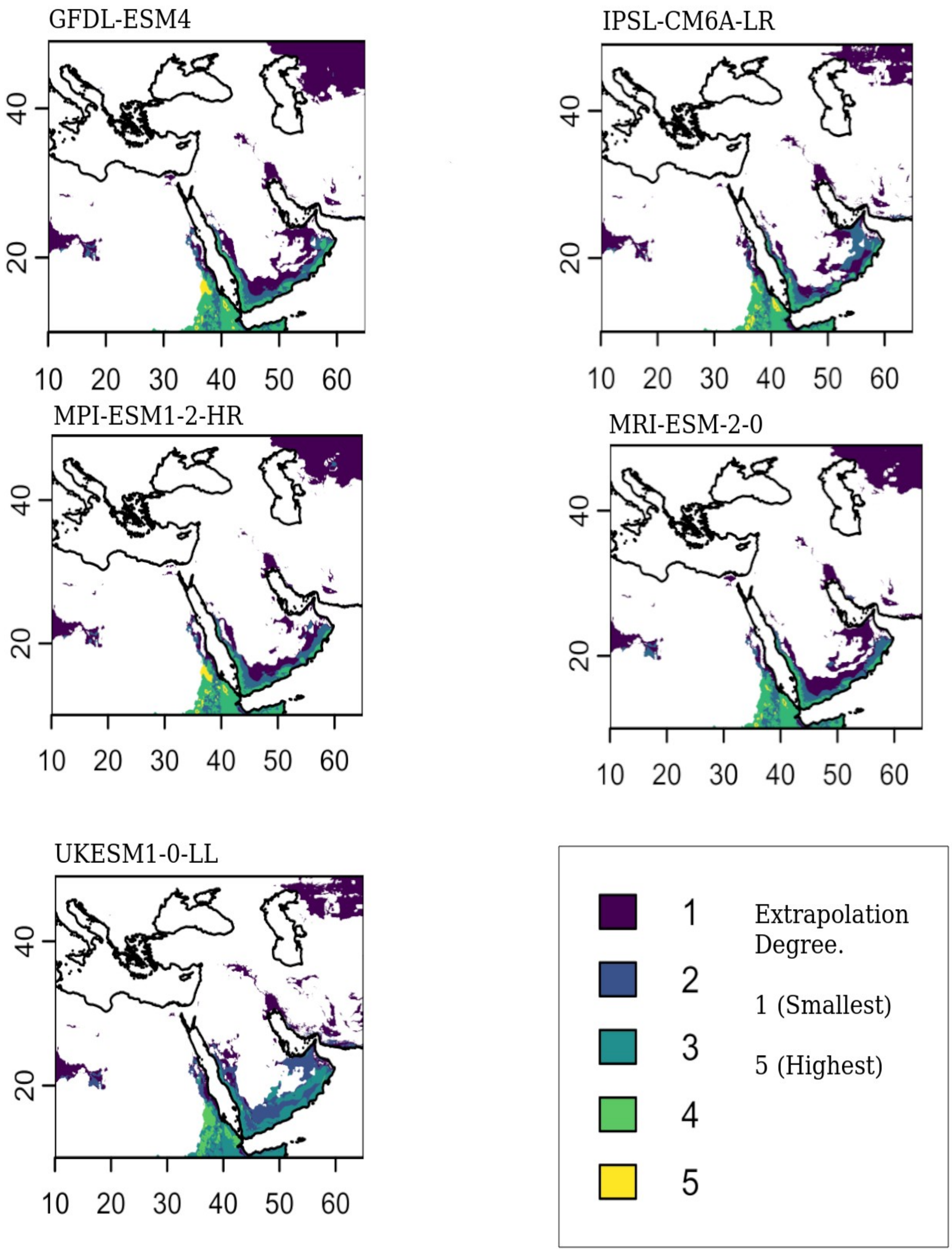


MOP (Mobility Oriented Pair Analysis) extrapolation maps for all SSP3-7.0 2071-2100 scenarios.

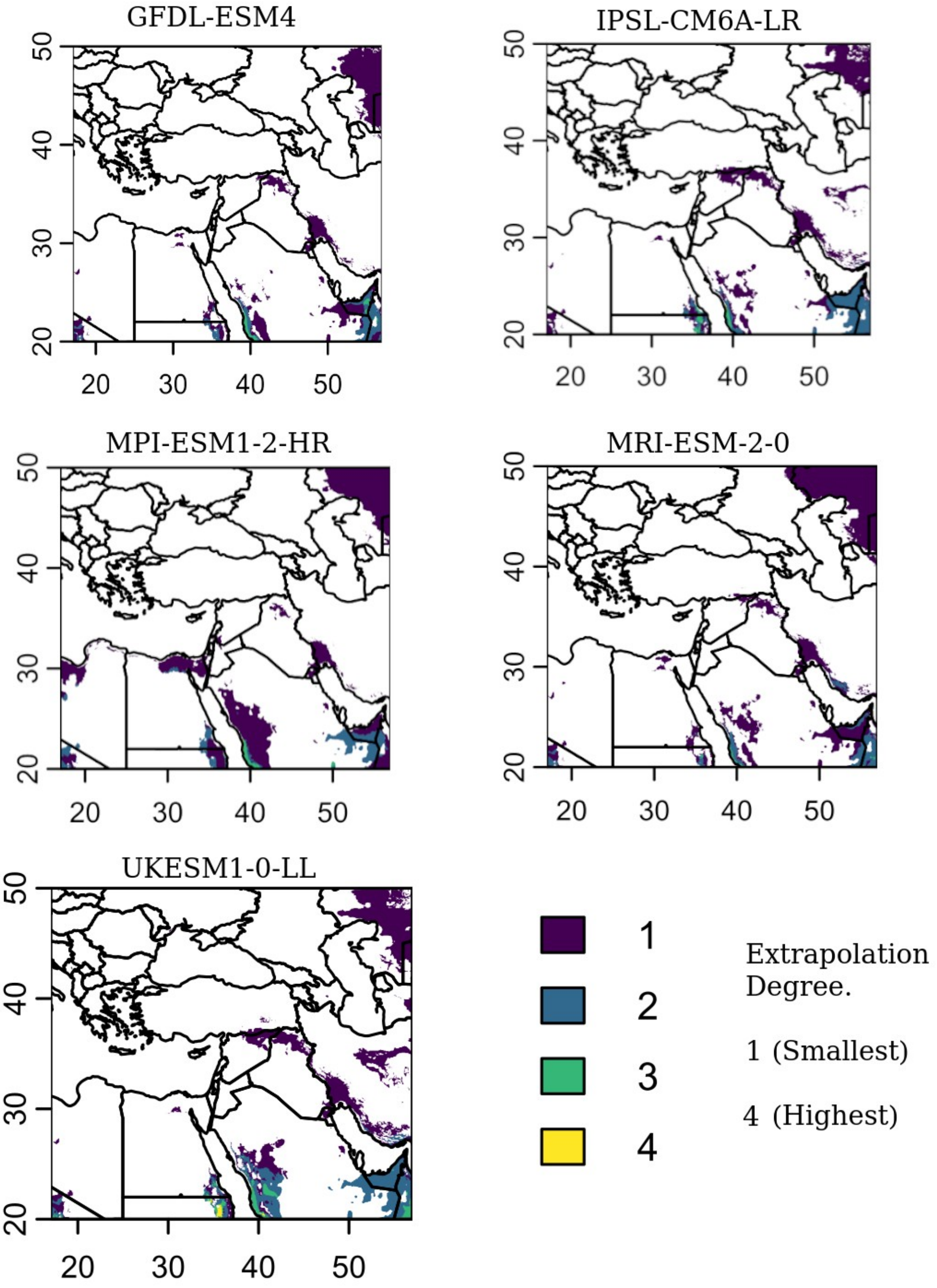


MOP (Mobility Oriented Pair Analysis) extrapolation maps for all SSP5-8.5 2071-2100 scenarios.

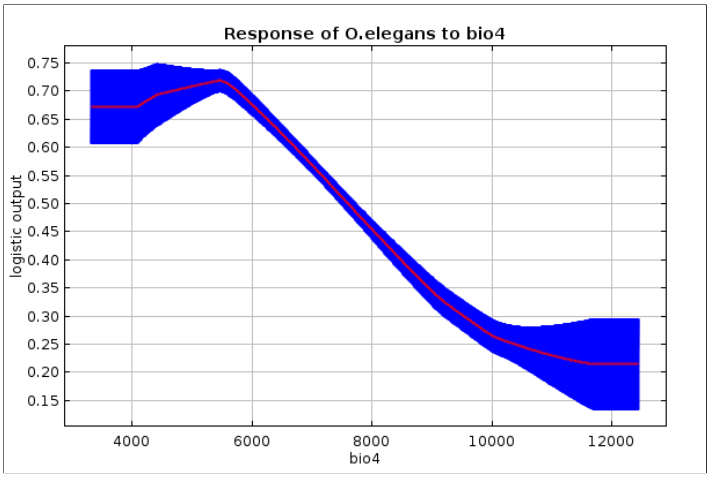


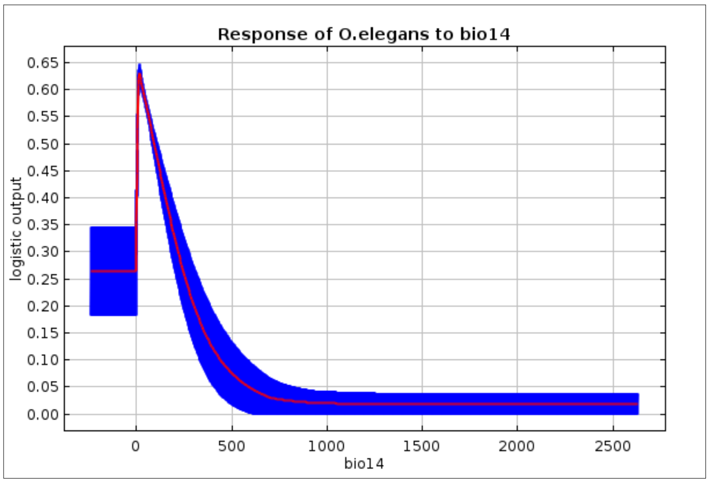


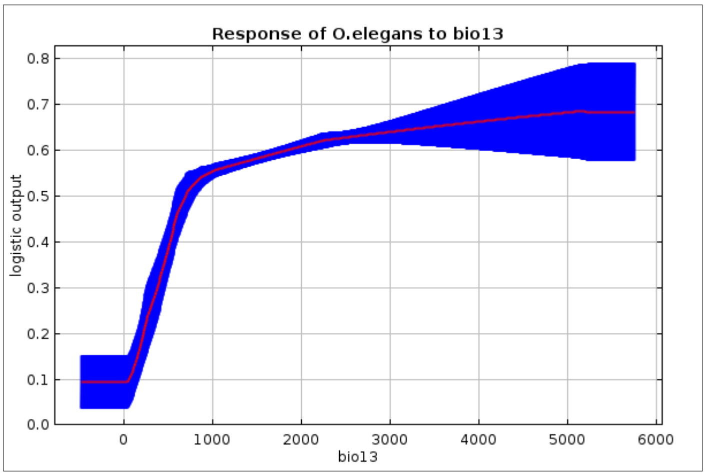


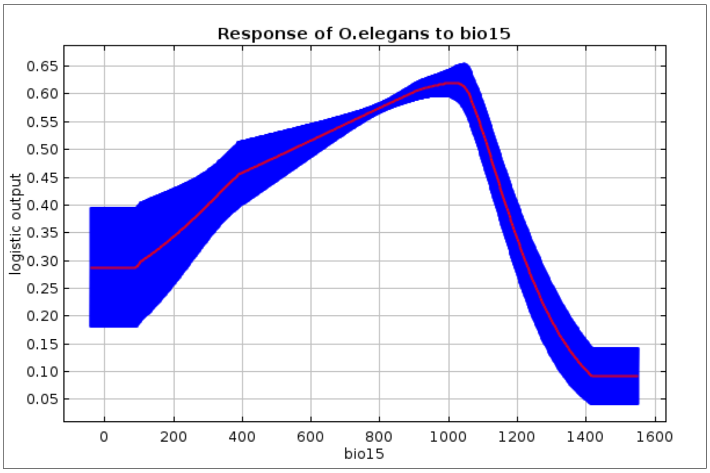


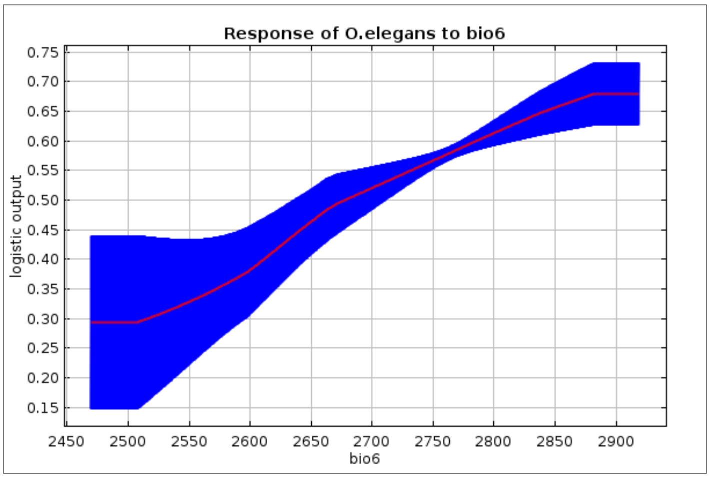


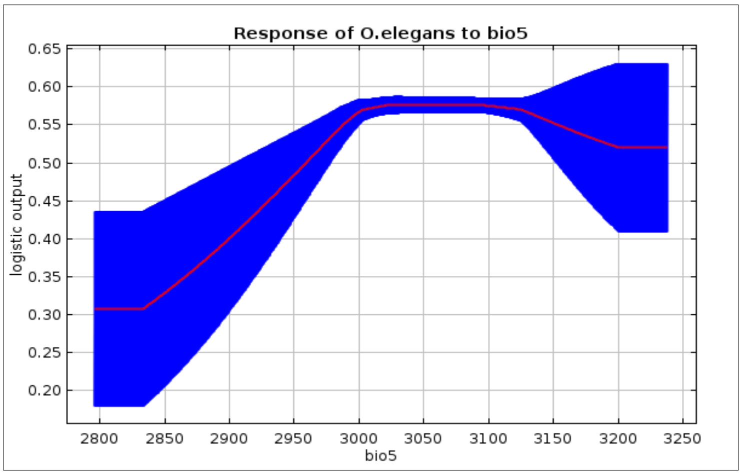


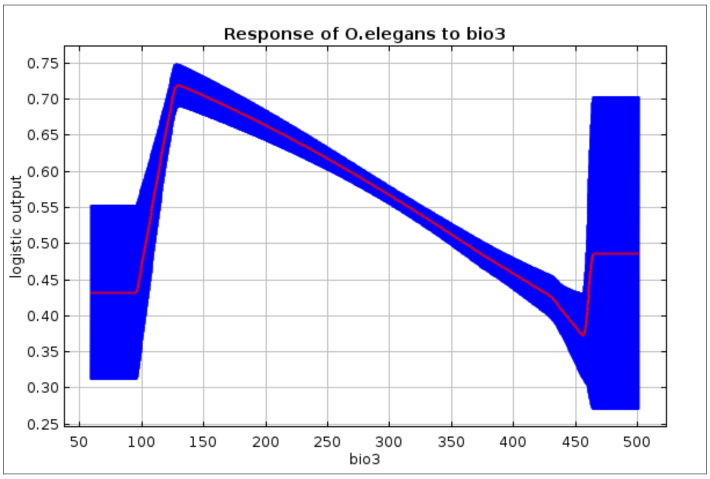


Response curves of all bioclimatic variables

For *O. elegans*
